# QTL mapping for seed morphology using the instance segmentation neural network in *Lactuca* spp

**DOI:** 10.1101/2022.05.19.492651

**Authors:** Kousuke Seki, Yosuke Toda

**Affiliations:** Nagano Vegetable and Ornamental Crops Experiment Station, 1066-1 Tokoo, Souga, Shiojiri, Nagano 399-6461, Japan; Phytometrics Co., Ltd., Shizuoka, Japan; Bioscience and Biotechnology Center, Nagoya University, Nagoya, 464-8602, Japan; Institute of Transformative Bio-Molecules (WPI-ITbM), Nagoya University, Chikusa, Nagoya 464-8602, Japan

**Author notes:** Corresponding Author: Kousuke Seki.

## Abstract

Wild species of lettuce (*Lactuca* sp.) are thought to have first been domesticated for oilseed contents to provide seed oil for human consumption. Although seed morphology is an important trait contributing to oilseed in lettuce, the underlying genetic mechanisms remain elusive. Since lettuce seeds are small, a manual phenotypic determination required for a genetic dissection of such traits is challenging. In this study, we built and applied an instance segmentation-based seed morphology quantification pipeline to measure traits in seeds generated from a cross between the domesticated oilseed type cultivar ‘Oilseed’ and the wild species ‘UenoyamaMaruba’ in an automated manner. Quantitative trait locus (QTL) mapping following ddRAD-seq revealed 11 QTLs linked to 7 seed traits (area, width, length, length-to-width ratio, eccentricity, perimeter length, and circularity). Remarkably, the three QTLs with the highest LOD scores, *qLWR-3*.*1, qECC-3*.*1*, and *qCIR-3*.*1*, for length-to-width ratio, eccentricity, and circularity, respectively, mapped to linkage group 3 (LG3) around 161.5 to 214.6 Mb, a region previously reported to be associated with domestication traits from wild species. These results suggest that the oilseed cultivar harbors genes acquired during domestication to control seed shape in this genomic region. This study also provides genetic evidence that domestication arose, at least in part, by selection for the oilseed type from wild species and demonstrates the effectiveness of image-based phenotyping to accelerate discoveries of the genetic basis for small morphological features such as seed size and shape.

## 1. Introduction

Lettuce (*Lactuca sativa* L.) cultivars can be classified into several types based on differences in plant morphology, such as crisphead, butterhead, romaine, leaf, latin, stem, and oilseed (Ryder 1999). Seed oil is also abundant in the seeds of wild lettuce species (Ramadan 1976). The oilseed type of lettuce was proposed to have been domesticated from wild species in ancient Egypt (Wei et al. 2021) and is considered to be the earliest domesticated type. In support of this notion, there are indications of the use of oilseed type lettuce from as early as ca. 2500 B.C. in Egypt (De Vries 1997). Lettuce cultivars from the oilseed type progress through the rosette stage very rapidly and bolt and flower early. However, the leaves are bitter and are thus not eaten as salad. The seeds are more round and larger than those of wild species, contain more vitamins soluble in seed oil, and were used for human consumption in Egypt.

For oilseed-type cultivars, seed morphology is an important agronomic trait that has been altered over the course of domestication. However, little is known about the genetic basis for seed morphology in lettuce, except for previous research focused on domestication traits based on quantitative trait locus (QTL) analysis (Hartman et al. 2013). One of the main limitations comes from the small size of lettuce seeds, making it difficult to evaluate seed morphology traits by hand. In seed crops such as soybean (*Glycine max*), rice (*Oryza sativa*), and wheat (*Triticum aestivum*), seed morphology is also an important trait related to crop yield in which phenotyping has been performed via several approaches (Tanabata et al. 2012; Williams et al. 2013; Gao et al. 2019; Kumawat and Xu 2021). For instance, seed dimensions have been measured manually with calipers or with image-processing software. These methods are labor-intensive and run the risk of bias due to human error. Automation is thus desirable for the precise phenotyping of seed morphology traits. Several software programs have been developed for plant phenotyping including seeds based on images captured with digital cameras, but the results are strongly influenced by the capture conditions. Indeed, overlapping seeds may be recognized as a single seed, resulting in an abnormal seed shape output. To address this issue, we implemented an image analysis pipeline for phenotyping morphological traits of lettuce seeds. Specifically, we trained a neural network with a synthetic dataset to identify and segregate the respective seed area from densely oriented seed images, regardless of their orientation (Toda et al. 2020).

The purpose of our study was to identify QTLs rapidly and reliably for seed morphology of oilseed-type cultivar by combining genotyping by double digest restriction-site associated DNA sequencing (ddRADseq) and phenotyping by automated image analysis. We also aimed to elucidate the genetic mechanisms related to domestication from wild species to oilseed-type cultivars. We characterized an F_2_ population derived from a cross between an oilseed-type cultivar and a wild species for QTL mapping of seed morphology.

## 2. Materials and methods

### 2.1. Plant materials

All plant materials were grown at the Nagano Vegetable and Ornamental Crops Experiment Station (Shiojiri City, Nagano prefecture, Japan; 36° 10′ N, 137° 93′ E). The lettuce cultivar ‘Oilseed’ derived from upper Egypt was obtained from the Centre for Genetic Resources, the Netherlands (CGN), under the stock number ‘CGN04769’. The wild *Lactuca* species ‘UenoyamaMaruba’ is the landrace of Nagano prefecture in Japan. A set of 173 F_2_ individuals derived from the ‘Oilseed’ × ‘UenoyamaMaruba’ cross was used for linkage analysis and to produce self-pollinated F_3_ lines that were used to evaluate the segregation patterns for genotypes and seed morphology of the F_2_ individuals. F_3_ seeds were spread onto white paper, and images were captured with a digital camera C5050Z (Olympus optical Co., Ltd, Japan) using transmission light. Each image size was 2560 × 1920 pixels at a resolution of 72 dpi.

### 2.2. Quantification of seed morphology by automated image analysis

An instance segmentation neural network (Mask R-CNN; He, K., et al., 2017) was trained and applied to extract morphological parameters of lettuce seeds, as described previously (Toda et al. 2020). Briefly, 28 single seed images of ‘UenoyamaMaruba’ were used to generate a synthetic image. The generated image resolution was 700 × 700 pixels, and each image was center-cropped to the size of 512 × 512 pixels for model training. Mask R-CNN was implemented on a Keras backend (https://github.com/matterport/Mask_RCNN) with default parameters. Since the number of images to be analyzed was limited (159 images), visual inspection against the inference result was used to decide whether the training was adequate. In our case, 40 epochs of training led to an acceptable result.

After isolating the seed area, morphological parameters of the respective regions were calculated using the measure.resionprops module of the sci-kit image library. Seed length (major_axis_length), width, (minor_axis_length), area, perimeter, eccentricity, and bounding box coordinate (bbox) were directly obtained from the above module (Table 1), and the metrics were calculated by the following formulas; length_to_width_ratio = major_axis_length/minor_axis_length; circularity = (4 × Pi × area) / perimeter^2^. However, the dataset may include data for incomplete seed shapes; e.g. when seeds are at the edge of the image, physically damaged, or partially covered and masked by other seeds. Therefore, such potential noise in the following analysis was excluded by the following criteria: bounding box coordinates that exceed a five-pixel margin from the edge and length_to_width_ratio that falls outside the 5% and 95% quantile range of the total population. The filtered dataset was normalized by a scaling factor of 43.833 (pixels/mm) before subsequent analysis.

**Table 1.**
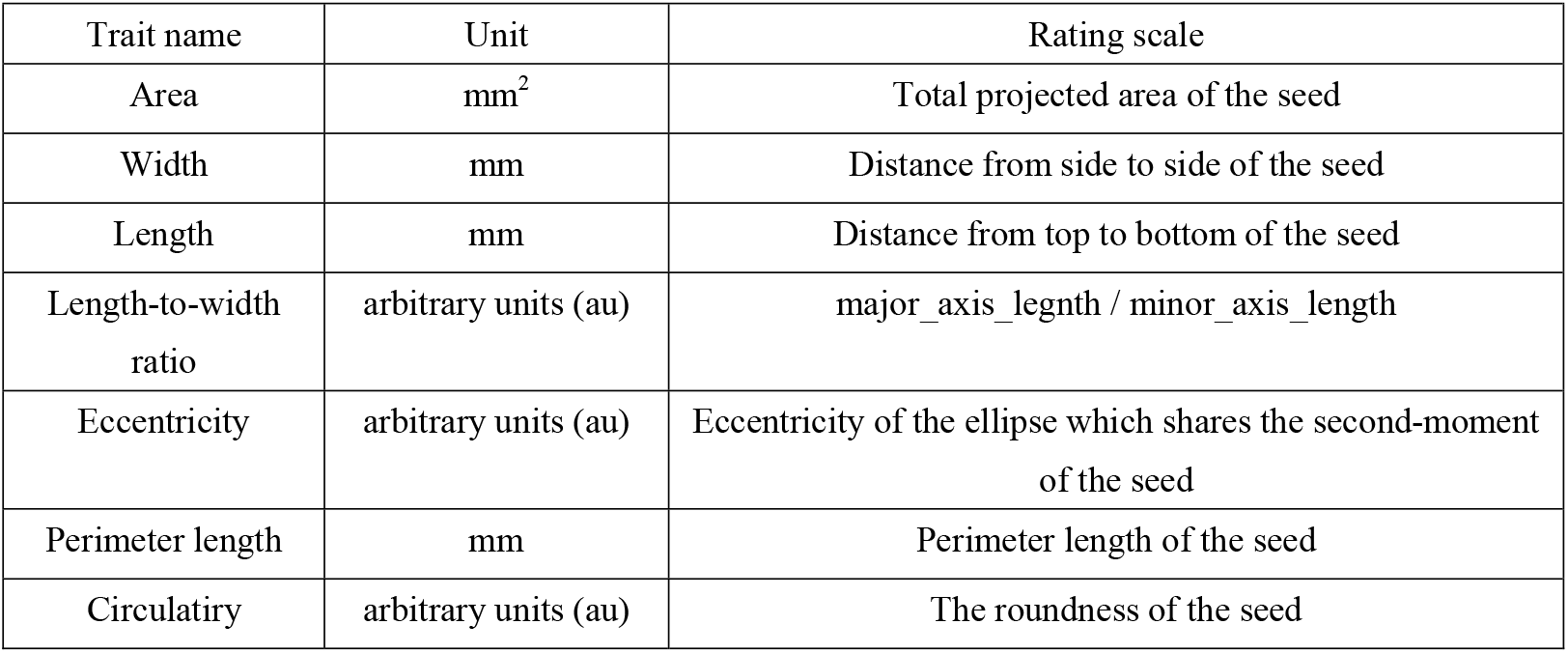
Traits measured on the parental cultivars and the F_2_ population derived from the ‘Oilseed’ × ‘UenoyamaMaruba’ cross. See Methods for details of the respective parameters.

### 2.3. Linkage map construction by ddRAD-seq analysis

Genomic DNA was extracted from leaves using the Nucleo-Spin Plant II Extract Kit (Machery-Nagel, Duren, Germany). ddRAD-seq library construction was performed following a previously described method (Matsumura et al. 2014; Seki et al. 2020). The ddRAD-seq libraries were sequenced on a HiSeqXten platform (Illumina, San Diego, CA, USA). Paired-end sequencing reads (150 bp × 2) were analyzed for ddRAD-seq tag extraction, counting, and linkage map construction using RAD-R scripts (Seki 2021). Reads were mapped with the RAD tags in each parent against the lettuce reference genome sequence (version8 from the crisphead cultivar ‘Salinas’, https://genomevolution.org/coge/GenomeInfo.pl?gid=28333). The linkage map was drawn using R/qtl (Broman et al. 2003). Raw sequence data (FASTQ) for this ddRAD-seq dataset were deposited in the DNA Data Bank of Japan (DDBJ) Sequence Read Archive (http://ddbj.nig.ac.jp/dra/index_e.html) under accession number DRA013652.

### 2.4. QTL detection by composite interval mapping (CIM)

QTL detection with CIM was conducted using the Haley–Knott regression of the R/qtl package in R (Broman et al. 2003). The genome-wide LOD threshold at the 1% and 5% significance levels was individually determined using a 10,000-permutation test for each trait. The proportion of phenotypic variance was calculated from the value at the peak, as indicated by CIM. A detailed script is described in CIM_script.R (https://github.com/KousukeSEKI/RAD-seq_scripts).

## 3. Results

### 3.1. Phenotyping seed morphology with instance segmentation

We applied our image analysis pipeline to phenotype seed morphology traits using seed images from 173 F_3_ lines and those of the two parental lines. Because we trained the neural network with a training set of seed images, our pipeline recognized overlapping seeds as individual seeds (Figure 1). To analyze seed morphology traits, we performed a number of post-processing steps: We removed outliers falling outside of a 12.5% (lower limit) and 87.5% (upper limit) quantile threshold for all traits for each individual. Across the generated dataset, we measured at least 71 seeds per line, with a maximum number of 399 seeds and a mean of 243. We applied a Student’s *t*-test to compare seed morphology traits between the two parents: We observed statistically significant differences in all seven traits (area, width, length, length-to-width ratio, eccentricity, perimeter length, and circularity) (Table 1, Table 2). A principal component analysis (PCA) using all data of seven traits from 173 F3 lines revealed that the first two principal components (PCs) explain 98.8% of the total standing variation (Figure 2). The length-to-width ratio, eccentricity, length, perimeter length, and circularity contributed to the first PC, with circularity being oriented in the direction opposite to that of length-to-width ratio and eccentricity. All traits contributed to the second PC. The direction of the length eigenvector was nearly identical to that of perimeter length. The direction of width differed from that of perimeter length and length. Notably, the area eigenvector was in between the direction of width and length. We calculated the Pearson’s correlation coefficients between the traits using all measurements of seed morphology to explore relationships. We observed a range of correlation values, from weak correlations to significant correlations reaching up to 0.99 (Table 3). In addition, Pearson’s correlation coefficients reflected the results of the PCA, with strong positive correlations between close eigenvectors (length-to-width ratio and eccentricity [0.99, *P* < 0.01], area and width [0.91, *P* < 0.01], between area and length [0.86, *P* < 0.01], and between area and perimeter length [0.92, *P* < 0.01]) and strong negative correlations oppositely pointing eigenvectors (length-to-width ratio and circularity [–0.96, *P* < 0.01]).

**Figure 1.**
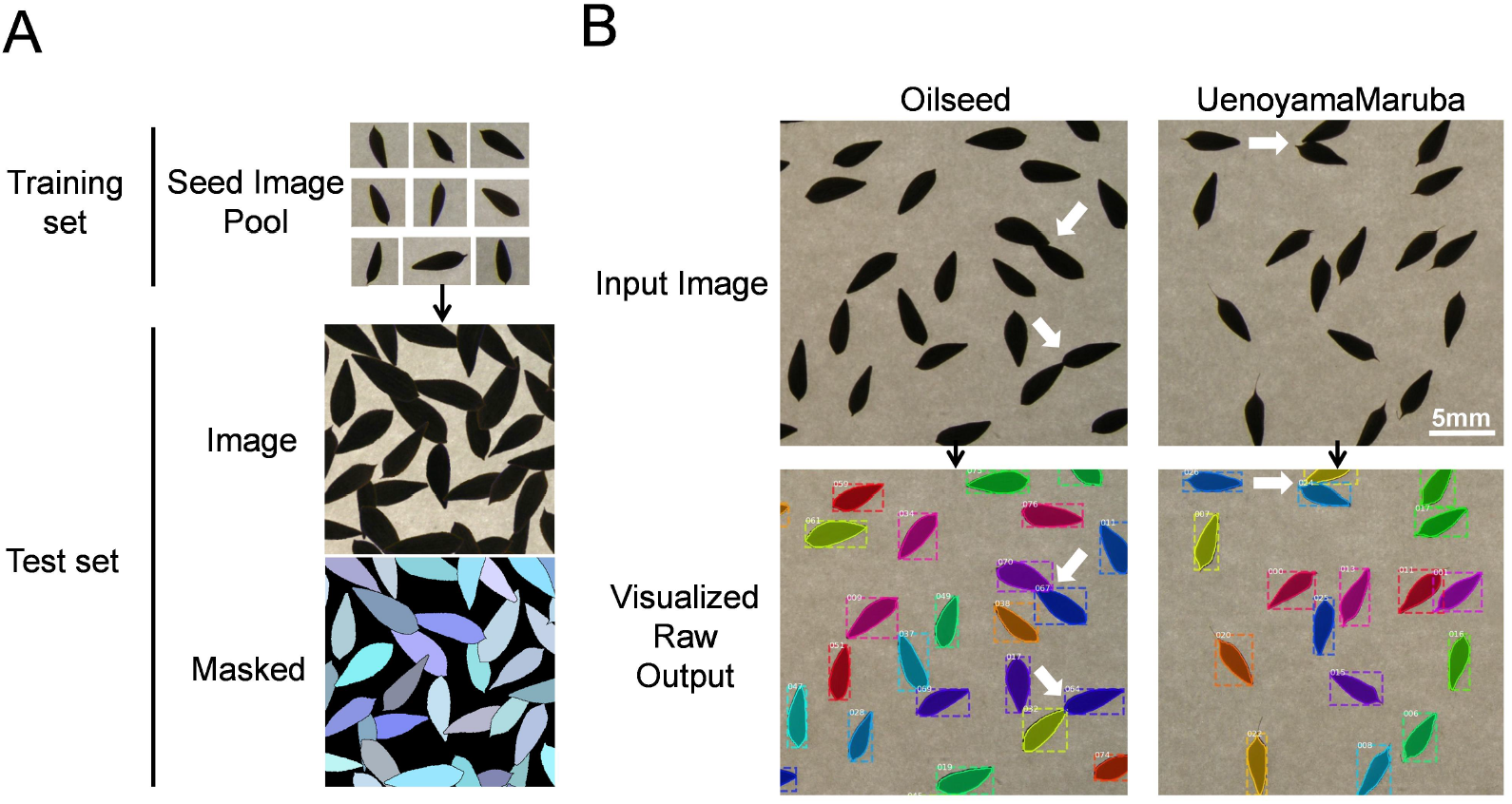
Overview of the image analysis pipeline using lettuce seeds. **a** Training pool with individual seed images to identify seeds and measure seed morphology traits. **b** Representative analyses of input images for the two parental cultivars. White arrows indicate overlapping seeds that are recognized as individual seeds by the image analysis pipeline.

**Table 2.** Phenotypic traits measured in the parental cultivars.

| Trait name | Oilseed |  |  |  | UenoyamaMaruba |  |  |  | *T*-test |
| --- | --- | --- | --- | --- | --- | --- | --- | --- | --- |
|  | Min | Mean | Max | SD | Min | Mean | Max | SD | *P*-value |
| Area | 2.52 | 3.51 | 4.14 | 0.39 | 2.91 | 3.02 | 3.15 | 0.08 | <0.01** |
| Width | 1.04 | 1.28 | 1.43 | 0.09 | 1.14 | 1.18 | 1.20 | 0.02 | <0.01** |
| Length | 3.00 | 3.54 | 3.85 | 0.20 | 3.27 | 3.36 | 3.47 | 0.06 | 0.003** |
| Length-to-Width ratio | 2.50 | 2.76 | 3.06 | 0.15 | 2.76 | 2.86 | 2.96 | 0.07 | 0.04* |
| Eccentricity | 0.92 | 0.93 | 0.95 | 0.01 | 0.93 | 0.94 | 0.94 | 0.003 | 0.03* |
| Perimeter length | 7.53 | 8.84 | 9.52 | 0.48 | 8.21 | 8.53 | 8.87 | 0.18 | 0.03* |
| Circulatiry | 0.52 | 0.56 | 0.60 | 0.02 | 0.50 | 0.52 | 0.55 | 0.01 | <0.01** |

\*\*, \* significant at the level of 1 % and 5 %, respectively

**Figure 2.**
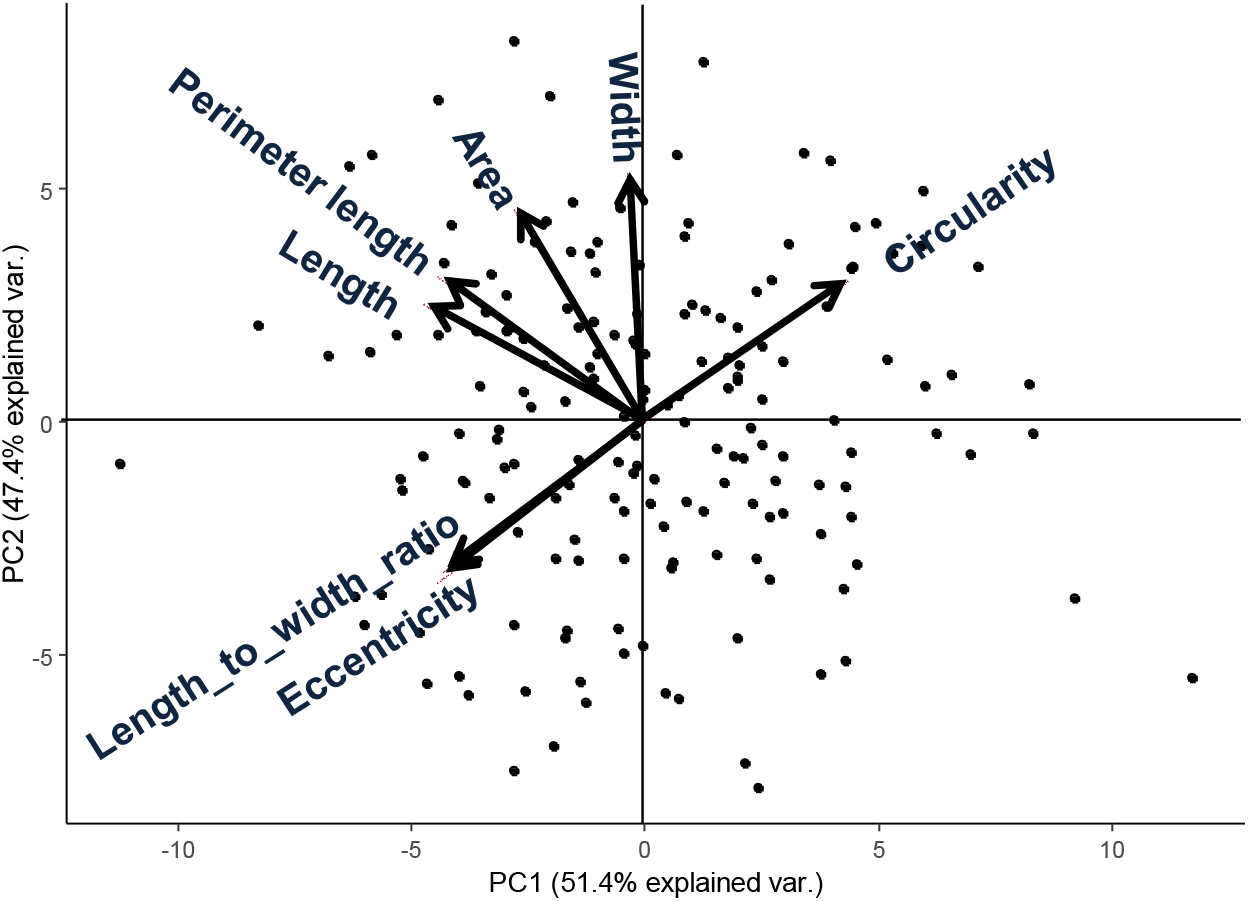
Principal component analysis from 173 F_3_ lines of lettuce seed morphological parameters. Arrows indicate eigenvectors of each descriptor.

**Table 3.**
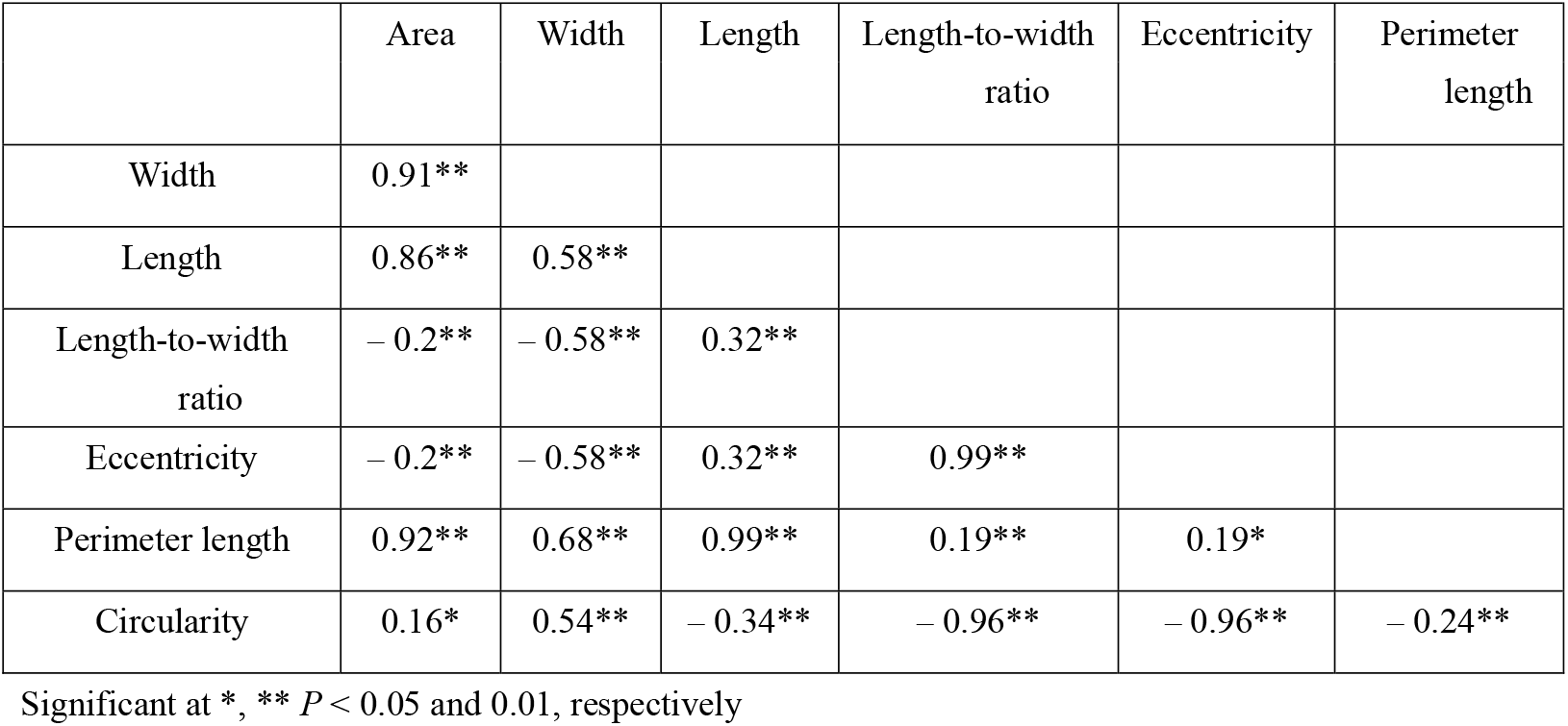
Pearson’s correlation coefficients of seed morphology traits using the phenotyping data of F_3_ seeds.

### 3.2. Genotyping by ddRAD-seq analysis

For genetic mapping of the loci controlling seed morphology, we conducted ddRAD-seq analysis for constructing a linkage map using an F_2_ population derived from a cross between the accessions ‘Oilseed’ and ‘UenoyamaMaruba’. Following sequencing of the ddRAD-seq libraries on an Illumina HiSeq instrument, we obtained 1,075,108 and 1,125,238 single reads (150 bp) for ‘Oilseed’ and ‘UenoyamaMaruba’, respectively. We converted the genotypes from ‘Oilseed’ and ‘UenoyamaMaruba’ genotypes to A and B, respectively. For the RAD-R scripts (Seki 2021), we employed mem as BWA mode, select as construction method, and 7US as correction approach. We then used the 1,870 pairs of RAD tags extracted from the two parents as codominant markers for genetic mapping of seed morphology for linkage map construction based on the genotypes of the 173 F_2_ individuals (Table 4). The total length of the resulting linkage map was 1,505.4 cM. The average inter-marker distance ranged from 0.5 (LG3) to 1.7 cM (LG9). The number of markers per linkage group ranged from 109 (LG9) to 371 (LG4).

**Table 4.** Summary of integrated lettuce linkage groups and genetic map information.

| Linkage groups | Total mapped tags | Common tags |  | Oilseed unique tags |  | UenoyamaMaruba unique tags |  | Linkage construct marker |  |  |
| --- | --- | --- | --- | --- | --- | --- | --- | --- | --- | --- |
|  |  |  |  |  |  |  |  | No. biallelic tags | Map length | Average interval between markers |
|  |  | No. RAD-tags | (%) | No. RAD-tags | (%) | No. RAD-tags | (%) |  | (cM) | (cM) |
| LG1 | 35396 | 3142 | 8.9 | 15556 | 43.9 | 16698 | 47.2 | 173 | 133.1 | 0.8 |
| LG2 | 35377 | 2971 | 8.4 | 16423 | 46.4 | 15983 | 45.2 | 214 | 140.6 | 0.7 |
| LG3 | 49415 | 7519 | 15.2 | 21610 | 43.7 | 20286 | 41.1 | 308 | 169.2 | 0.5 |
| LG4 | 67093 | 8632 | 12.9 | 29791 | 44.4 | 28670 | 42.7 | 371 | 236.1 | 0.6 |
| LG5 | 64605 | 12409 | 19.2 | 26698 | 41.3 | 25498 | 39.5 | 227 | 203.1 | 0.9 |
| LG6 | 42377 | 11312 | 26.7 | 15830 | 37.4 | 15235 | 36.0 | 127 | 134.1 | 1.1 |
| LG7 | 46046 | 16001 | 34.8 | 15006 | 32.6 | 15039 | 32.7 | 120 | 122.2 | 1.0 |
| LG8 | 64692 | 15570 | 24.1 | 25376 | 39.2 | 23746 | 36.7 | 221 | 179.6 | 0.8 |
| LG9 | 51409 | 19153 | 37.3 | 16593 | 32.3 | 15663 | 30.5 | 109 | 187.4 | 1.7 |
| Total | 456410 | 96709 | 21.2 | 182883 | 40.1 | 176818 | 38.7 | 1870 | 1505.4 | 0.8 |

### 3.3. QTL mapping of seed morphology

We used composite interval mapping (CIM) to detect QTLs, using the seed morphology trait values determined by the pipeline and the genotype data from the ddRAD-seq across 173 F_2_ individuals. We detected 11 QTLs for 7 traits (Table 5). Importantly, the QTLs with the highest logarithm of odds (LOD) scores, *qLWR-3*.*1, qECC-3*.*1*, and *qCIR-3*.*1*, associated with the length-to-width ratio, eccentricity, and circularity, respectively, mapped to LG3 between 161.5 and 214.6 Mb. These three QTLs each accounted for approximately 27.95%, 28.55%, and 29.84% of the phenotypic variation explained (PVE) for their respective trait. We observed significant differences in the phenotypic values of putative homozygote and heterozygote individuals for each of the three traits (Figure 3), suggesting that each trait is controlled by one semi-dominant locus. We also detected two QTLs linked to both length and perimeter length at almost the same position on LG5 and LG8. The QTLs *qWID-7*.*1* and *qWID-8*.*1* linked to width mapped to LG7 and LG8, while *qARE-8*.*1* associated with area mapped almost to the same position as *qWID-8*.*1* (Table 5).

**Table 5.** QTL detected by composite interval mapping in the F_2_ population derived from the ‘Oilseed’ × ‘UenoyamaMaruba’ cross.

| QTL name | Trait name | Linkage group | Marker | Genetic distance (cM) | Physical position (cM) | Threshold LOD (1%) | Threshold LOD (5%) | LOD | Additive effect | Dominant effect | $R^2$ (%) |
| --- | --- | --- | --- | --- | --- | --- | --- | --- | --- | --- | --- |
| *qLWR-3.1* | Length-to-Width ratio | 3 | LG3_v8_176.7Mbp to LG3_v8_214.6Mbp | 16.88 | 138.0 | 4.57 | 3.88 | 12.46 | 0.1628 | 0.0167 | 27.95 |
| *qECC-3.1* | Eccentricity | 3 | LG3_v8_176.7Mbp to LG3_v8_214.6Mbp | 16.88 | 138.0 | 4.77 | 4.00 | 12.77 | 0.0051 | 0.0006 | 28.55 |
| *qCIR-3.1* | Circularity | 3 | LG3_v8_161.5Mbp to LG3_v8_189.0Mbp | 14.11 | 133.0 | 4.63 | 3.97 | 12.62 | -0.0191 | -0.0024 | 29.84 |
| *qCIR-4.1* | Circularity | 4 | LG4_v8_73.0Mbp to LG4_v8_73.4Mbp | 1.81 | 44.6 | 4.63 | 3.97 | 4.01 | 0.0032 | 0.0126 | 7.35 |
| *qLEN-5.1* | Length | 5 | LG5_v8_230.7Mbp to LG5_v8_260.7Mbp | 19.43 | 141.0 | 4.62 | 3.88 | 4.58 | -0.0858 | -0.0399 | 10.03 |
| *qLEN-8.1* | Length | 8 | LG8_v8_293.1Mbp to LG8_v8_307.9Mbp | 11.88 | 172.1 | 4.62 | 3.88 | 4.66 | 0.0070 | -0.1280 | 10.49 |
| *qPER-5.1* | Perimeter length | 5 | LG5_v8_253.7Mbp to LG5_v8_260.7Mbp | 6.27 | 141.0 | 4.79 | 4.01 | 4.27 | -0.1958 | -0.1051 | 9.30 |
| *qPER-8.1* | Perimeter length | 8 | LG8_v8_298.8Mbp to LG8_v8_305.4Mbp | 3.46 | 172.1 | 4.79 | 4.01 | 5.01 | 0.0092 | -0.3185 | 11.24 |
| *qWID-7.1* | Width | 7 | LG7_v8_165.9Mbp to LG7_v8_192.4Mbp | 20.64 | 106.5 | 4.37 | 3.78 | 4.54 | -0.0336 | 0.0077 | 10.88 |
| *qWID-8.1* | Width | 8 | LG8_v8_279.1Mbp to LG8_v8_293.1Mbp | 4.01 | 166.0 | 4.37 | 3.78 | 3.80 | -0.0258 | -0.0205 | 8.44 |
| *qARE-8.1* | Area | 8 | LG8_v8_279.1Mbp to LG8_v8_295.3Mbp | 5.70 | 163.1 | 4.67 | 3.95 | 4.52 | -0.0703 | -0.1623 | 11.21 |

**Figure 3.**
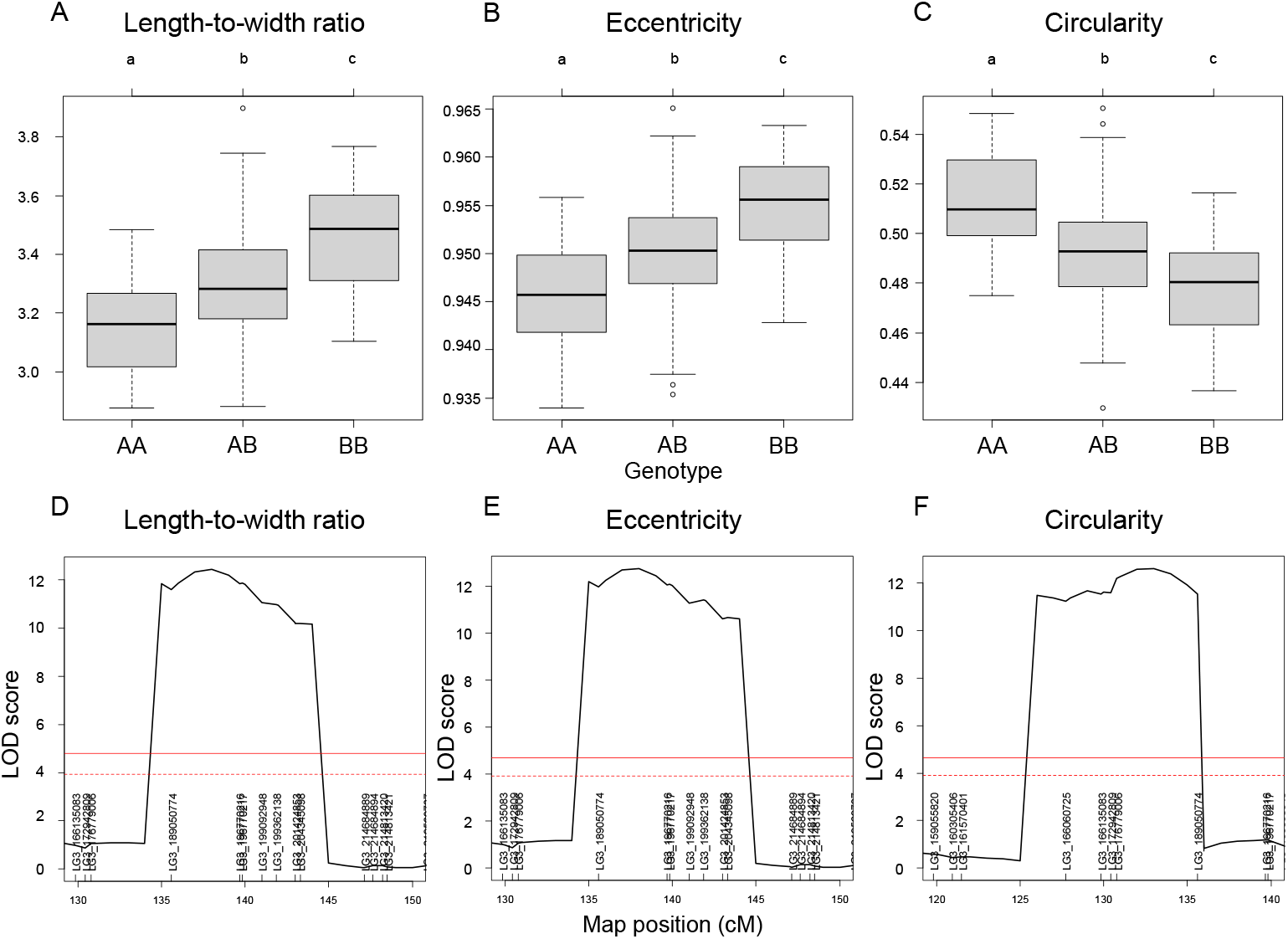
QTL mapping results for selected seed morphology traits **a-c** Phenotypic difference for the traits length-to-width ratio (a), eccentricity (b), and circularity (c) between the three possible genotypes at three major QTLs with high LOD scores. **d-f** QTL peaks with LOD scores at each locus. Different lowercase letters represent significant differences (*P* < 0.01).

## 4. Discussion

Lettuce, a diploid species (2n = 2x = 18), is a self-pollinated crop with a large genome (∼ 2.3 Gb) (Reyes-Chin-Wo et al. 2017). Cultivated lettuce exhibits a low rate of intraspecific polymorphisms (Truco et al. 2007; Simko and Hu 2008). In this study, we produced a segregating F_2_ population derived from a cross between an oilseed-type cultivar and a wild species of lettuce to maximize the detection of polymorphisms. We thus developed 1,870 RAD markers and assigned them to 9 linkage groups by ddRAD-seq rapidly (Table 4). While our previous report detected the rate of common tags between parents of crisphead type to be as high as 60% (Seki et al. 2020), the two parental lines used here only shared 21.2% of common tags. This result suggests that polymorphisms between oilseed type and wild lettuce species are more numerous than between crisphead-type lettuce cultivars. With the help of our image analysis pipeline, we obtained phenotypic data for 42,036 seeds quickly and with high performance, which would be impossible to easily accomplish manually. The pipeline successfully recognized overlapping seeds and separated them into individual seeds, which is instrumental in large-scale phenotyping without having to place seeds evenly apart manually. The data collected with our pipeline correctly identified the bigger and rounder seeds characteristic of oilseed-type lettuce compared to the wild species (Figure 1, Table 2). A more rounded seed shape would be advantageous to accumulate more seed oil. Perhaps the initial artificial selection during domestication was for uniform germination, for which seed morphology and size are important determinants. Using the genotyping data generated by ddRAD-seq and the above phenotyping data, we detected 11 QTLs for the 7 seed morphology traits (Table 5). Of these, three QTLs, *qLWR-3*.*1, qECC-3*.*1*, and *qCIR-3*.*1*, exhibited major effects on their respective trait (PVE > 25%). PCA illustrated the relationships between traits using the phenotyping data gathered from F_3_ seeds (Figures 3 and 4). We measured a strong and positive correlation (R = 0.99) between perimeter length and seed length, while these two parameters were less strongly correlated with seed width (R = 0.68 and 0.58, respectively), reflecting the greater influence of seed length over seed width on seed perimeter length. In support of this observation, we detected two QTLs mapping to the same genomic regions on LG5 and LG8 for each of the two traits (Table 5). Similarly, seed area appeared to be positively strongly correlated with seed width (R = 0.91), but not seed length (R = 0.86). We detected a common QTL associated with seed area and width on LG8 (Table 5), in the same location as a previously reported QTL for seed width (Hartman et al. 2013). The length-to-width ratio, eccentricity, and circularity associated with the characteristic round shape features of oilseed-type lettuce also displayed very strong correlations among them, prompting us to take a closer look at their underlying genetic basis (Table 3). Remarkably, the major QTLs associated with these three traits mapped to almost the same region on LG3, from 161.5 to 214.6 Mb (Figure 3). We hypothesize that this genomic region harbors a key causal locus associated with the domestication of seed morphology from wild-type to oilseed type lettuce. Domestication involves the genetic modification of wild species due to selection for a given phenotype that is beneficial to humans and is often determined by few loci with major effects (Abbo et al. 2012). Although it is difficult to precisely compare genetic maps because of the limited number of markers, the same regions on LG3 and LG7 were reported to be associated with clusters of QTLs linked with domestication for the traits seed output, leaf shape, and late flowering in lettuce (Hartman et al. 2013). Thus, our hypothesis that the genomic region on LG3 mainly reflects the domestication of seed morphology traits agrees with the results of previous genetic studies.

Due to the recent advances in computation power, approaches based on deep learning are becoming accessible to agriculture (de Medeiros et al. 2021; Nehoshtan et al. 2021). Our data demonstrate that deep learning–based approaches could significantly contribute to developing the field of not only plant science but also agriculture for plant phenotyping.

## Author contributions

KS and YK planned the experiments. KS developed the mapping population, performed ddRAD-seq, and phenotyping data analysis. YK performed crop seed phenotyping. KS and YK wrote and approved the manuscript.

## Acknowledgements

This work was supported in part by a Grant-in-Aid for Scientific Research on Innovative Areas (21H05152 for YT). We thank Dr. Ken Naito and all the organizers for hosting “Society of Post Youth Agronomists (SPY-A)”, a research conference which not only provided the opportunity of the coauthors’ acquaintance but also led to the collaboration of this research.

